# A Myelin Map of Trunk Folds in the Elephant Trigeminal Nucleus

**DOI:** 10.1101/2023.11.15.567239

**Authors:** Noémie Reveyaz, Undine Schneeweiß, Olivia Heise, Ben Gerhardt, Andreea M. Gui, Lena V. Kaufmann, Jette Alfken, Jakob Reichmann, Tim Salditt, Thomas Hildebrandt, Michael Brecht

## Abstract

Elephants have elaborate trunk skills and large, but poorly understood brains. Here we study trunk representations in elephant trigeminal nuclei, which form large protrusions on the ventral brainstem. These ventral brainstem protrusions have previously been referred to as inferior olive, but a delineation of the olivo-cerebellar tract reveals these (trigeminal) nuclei are not connected to the cerebellum via climbing fibers. In contrast, the olivo-cerebellar tract connects to a large dorsolateral nucleus with a serrated cellular architecture, the putative elephant inferior olive. Dense vascularization and intense cytochrome-oxidase reactivity distinguish several elongated trigeminal putative trunk modules, which repeat in the anterior-posterior direction. We focus on the most anterior and largest of these units, the putative nucleus principalis trunk module. Module neuron density is low and non-neural cells outnumber neurons by ∼108:1. Dendritic trees are elongated along the axis of axon bundles (myelin stripes) transversing the trunk module. Synchrotron X-ray-phase-contrast tomography suggests myelin-stripe-axons transverse the trunk module. We show a remarkable correspondence of trunk module myelin stripes and trunk folds. Myelin stripes show little relation to trigeminal neurons and stripe-axons appear to often go nowhere; we suggest that myelin stripes might serve to separate trunk-fold domains rather than to connect neurons. Myelin-stripes-to-folds mapping allowed to determine neural magnification factors, which changed from 1000:1 proximally to 5:1 in the trunk finger. Asian elephants have fewer (∼640,000) trunk-module neurons than Africans (∼740,000) and show enlarged representations of trunk parts involved in object wrapping. The elephant trigeminal trunk module is exquisitely organized into trunk-fold-related units.

## Introduction

Elephants are the largest extant terrestrial animals and rely on their trunks to acquire huge amounts of food. The trunk is a fusion organ of the nose and upper lip. Nose lip fusion occurs in the fourth month of elephant fetal development (Fischer & Trautmann, 1987; Schulz et al., 2023). The trunk is an immensely muscular structure (Cuvier, 1796; Cuvier and Dumeril, 1838; Shoshani, 1982) that contains about 90,000 muscle fascicles (Longren et al., 2023). Not surprisingly, the trunk’s prime motor control structure, the facial nucleus, is very large (Maseko et al., 2013) and shows an elaborate cellular architecture (Kaufmann et al., 2022). Trunks have prominent folds that differ between African (*Loxodonta africana*) and Asian (*Elephas maximus*) elephants (Schulz et al., in prep). Interestingly, the object-grasping behavior of African and Asian elephants differs considerably. Asian elephants have a single trunk finger and tend to wrap their trunk around objects, whereas African elephants prefer pinching objects with their two trunk fingers (Racine, 1980). Elephants are skillful with their trunks (Shoshani, 1992), have a high tactile sensitivity (Dehnhardt, Friese, and Sachser, 1997), and even acquire dexterous manipulative behaviors such as banana peeling (Kaufmann et al., 2023). Tactile feedback is of great significance for trunk behaviors because elephants have only limited visual abilities. Recently, Deiringer et al. (2023) investigated trunk whiskers and observed use-dependent whisker lateralization, dense whisker arrays on the trunk tip and the ventral trunk, and marked whisker differences between African and Asian elephants. The sensory periphery of elephant trunks was investigated in a landmark study by Rasmussen & Munger (1996), who described dense innervation patterns in the elephant fingertip. The whole elephant trunk is massively innervated by large trigeminal ganglia (Sprinz, 1952; Purkart et al., 2022). The cerebellum of elephants is very large both in relative and absolute terms (Maseko et al., 2012) and it is believed to be an important part of the control of the trunk movement, as a sensory and motor processing area (Maseko et al., (2013). The turning point in the investigation of the mammalian trigeminal system was the description of the cortical whisker barrels by Woolsey & Van der Loss (1970 and this work informed our approach to the elephant brainstem. The recognition of the cortical barrel pattern has led to thousands of follow-up studies, which included the discovery of whisker-related thalamic so-called barreloids (Van der Loos, 1976) and whisker-related units in the trigeminal brainstem, so-called barrelettes (Belford & Killackey, 1979; Ma, 1991). As shown by Belford & Killackey (1979) and Ma (1991) the brainstem contains several topographic trigeminal representations, which repeat in anterior to posterior direction. Specifically, these studies identified the most anterior one as the largest sensory trigeminal representation (the nucleus principalis) and several smaller more posterior trigeminal sensory nuclei. We will adopt the trigeminal terminology established by these authors (Belford & Killackey, 1979; Ma, 1991).

A variety of excellent studies have investigated the cellular statistics of elephant brains and indicated elephant brains are not simply scaled-up mouse brains. Specifically, investigators found much lower neuronal densities in elephant brains than in rodents (Haug, 1987; Herculano-Houzel et al., 2014). A prominent difference between small and large brains is the increased amounts of white matter in larger brains. Myelin sheaths, which give white matter its whitish shine, and the enwrapped axons are usually thought to form a supply and connectivity system, an idea we will question in our study.

We pursued the following questions: (i) Can candidate regions for the elephant trigeminal trunk representation be identified? (ii) If yes, can multiple sensory trigeminal nuclei be identified as in other mammals? (iii) What is the neuroanatomical structure of the elephant brainstem trunk representations? (iv) Do the elaborate myelin structures in the elephant trigeminal nuclei form an axonal supply system? (v) How does the organization of elephant brainstem trunk representations relate to the differential trunk morphology and grasping behavior of African and Asian elephants? We tentatively identified an elephant brainstem trunk module characterized by intense metabolism and vascularization. The putative trunk module contains a myelin map of trunk folds. This myelin map allows precise mapping of the neural topography of the trunk representation and reveals species differences between African and Asian elephants.

## Results

### Overview

Determining trigeminal representations in elephants is challenging because invasive recordings or invasive viral tracing methods cannot be applied. We proceeded to build a hypothesis on the elephant trigeminal brainstem trunk in seven steps. First, we identified a candidate module for the brainstem trunk representation. Second, we used peripherin-antibody staining to delineate the elephant olivo-cerebellar tract. This analysis indicated to us the ventral brainstem bumps do not correspond to the elephant inferior olive as previously thought (Maseko et al. 2013). Third, we showed that this module architecture repeats in the anterior-posterior direction in the elephant brainstem. Fourth, we characterized the cellular organization of this putative trunk module. Fifth, we documented a close correspondence between the myeloarchitecture of this module and the folds of the elephant trunk. Sixth, we applied synchrotron X-ray tomography to assess the microscopic architecture of myelin stripes. Seventh, we showed that species-specific differences in trunk structure have correlates in the putative trunk module.

### A metabolically highly active, strongly vascularized putative trunk module

The brain of the Asian elephant cow Burma is shown in Figure 1A. In rodents, the sensory trigeminal nuclei are observed posterior to the pons. As shown in Figure 1B, in a ventral view of the brain stem of Burma, about 1 cm posterior to the (large) pons, a pair of large bumps is obvious on the ventral brainstem surface. By size and pronounced protrusion, these bumps on the ventral brainstem distinguish elephant brains from those of other mammals. Much smaller bumps are seen in a similar position in the human brain, where they contain the inferior olive. Accordingly, the bumps of the elephant brain have been referred to as the elephant inferior olive (Shoshani et al., 2006; Maseko et al., 2013; Verhaart and Kramer, 1958; Verhaart 1962), but our investigation did not support this idea. We sectioned the elephant medulla and stained sections for cytochrome oxidase reactivity, a mitochondrial enzyme, the activity of which is closely related to tissue energy consumption. Trigeminal nuclei tend to show intense activity in cytochrome oxidase reactivity (Belford & Killackey, 1979; Ma, 1991). A cytochrome oxidase-stained coronal section through the bump of the Asian elephant bull Raj is shown in Figure 1C. We found that the bump contained the most intense cytochrome oxidase reactivity in the elephant brainstem and (to the extent that we performed such staining in other brain regions) the rest of the elephant brain. Three cytochrome-reactive modules (a putative trunk module, a putative nostril module, and a putative lower lip/jaw) are obvious, the largest of which we refer to as a putative trunk module (Figure 1D). The putative trunk module is elongated and we hypothesize that a particularly intensely cytochrome oxidase reactive region at the ventrolateral pole of the module corresponds to the dorsal finger representation (Figure 1C, D). We provide a detailed justification for our assignments of a putative trunk module, a putative nostril module, and a putative lower lip/jaw trigeminal module in Figure 2. We also studied brain stem sections in the African elephant (Figure 1E) and identified a similar putative trunk module there. We investigated the vascularization of this module, which was evident from the cytochrome oxidase reactivity of erythrocytes in blood vessels of our non-perfused elephant brains (Figure 1F), and found that the trunk module stands out from the rest of the brainstem (Figure 1G). Specifically, it contains about twice as many blood vessels per volume as the remainder of the brainstem (Figure 1H). In parasagittal sections, the putative trunk module had a compact appearance much like the trigeminal nuclei of other mammals (Figure 1I). In parasagittal sections lateral to the putative trunk module we observed a nucleus with a very distinct banded cellular appearance (Figure 1J), a cellular architecture characteristic of the inferior olive of other mammals (Brodal et al. 1980). We conclude that the elephant brainstem contains a large, highly vascularized, and highly cytochrome-oxidase reactive elongated putative trunk module.

**Figure 1.**
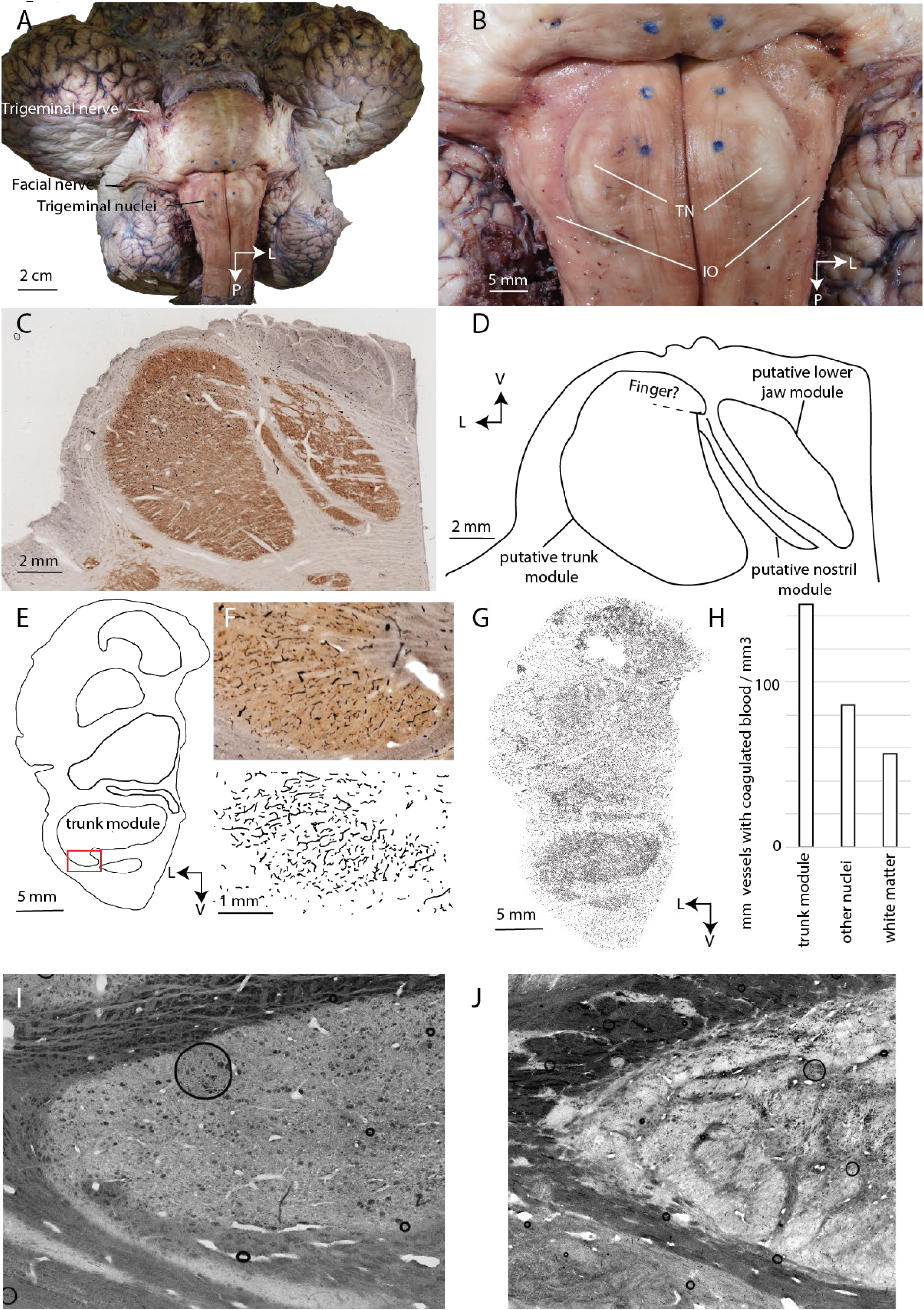
Appearance, metabolism, blood supply, and identification of the elephant trigeminal nuclei. **A,** ventral view of the brain of Burma, a 52-year-old female Asian elephant. **B**, ventral view of the brainstem of Burma. Note the large pons (upper part of the photograph). Posterior to the pons a pair of prominent bumps are visible, the putative trigeminal nuclei (TN) of the elephant. The trigeminal nuclei bumps are observed, wherein in humans and other mammals the inferior olive is found and was referred to as inferior olive by previous authors (Shoshani et al., 2006; Maseko et al., 2013; Rasenberger, 2019). The inferior olive can be identified at an unusual lateral position in elephants, however (IO). **C**, 60 µm coronal section through the trigeminal nuclei of Raj, a four-year-old elephant bull, stained for cytochrome-oxidase reactivity, a mitochondrial enzyme, the reactivity of which reflects constitutive metabolic activity. The trigeminal nuclei show some of the strongest cytochrome-oxidase reactivity (indicated by the brown color) in the elephant brain and strong cytochrome-oxidase reactivity is typical for the trigeminal nuclei of many mammals. **D**, a drawing of putative trigeminal subnuclei stained in **C**. Note the compact shape of the trunk module, which is unlike the inferior olive of mammals. **E**, drawing of a coronal section stained for cytochrome-oxidase reactivity through the brainstem of Indra a female African elephant, borders of nuclei are outlined. **F**, upper, micrograph of cytochrome-oxidase reactivity in the putative trunk fingertip representation. Lower, drawing of cytochrome-oxidase reactive erythrocytes in blood vessels. Note the high density of vessels inside of the fingertip but not in the surrounding tissue. **G**, drawing of the entire brainstem section. The putative trunk module stands out from the rest of the brainstem in terms of vessel density. **H**, quantification of blood vessel length in various parts of the brainstem. Note that not all vessels contain erythrocytes and are stained, i.e., measures of blood vessel length are lower bound estimates. **I**, Brightfield micrograph of a parasaggital section through the trigeminal trunk module. Note the compact cellular architecture. **J**, Brightfield micrograph of a parasaggital section through the inferior olive. Note the banded cellular architecture that is characteristic of the mammalian inferior olive. L = lateral; V = ventral; P = posterior.

**Figure 2.**
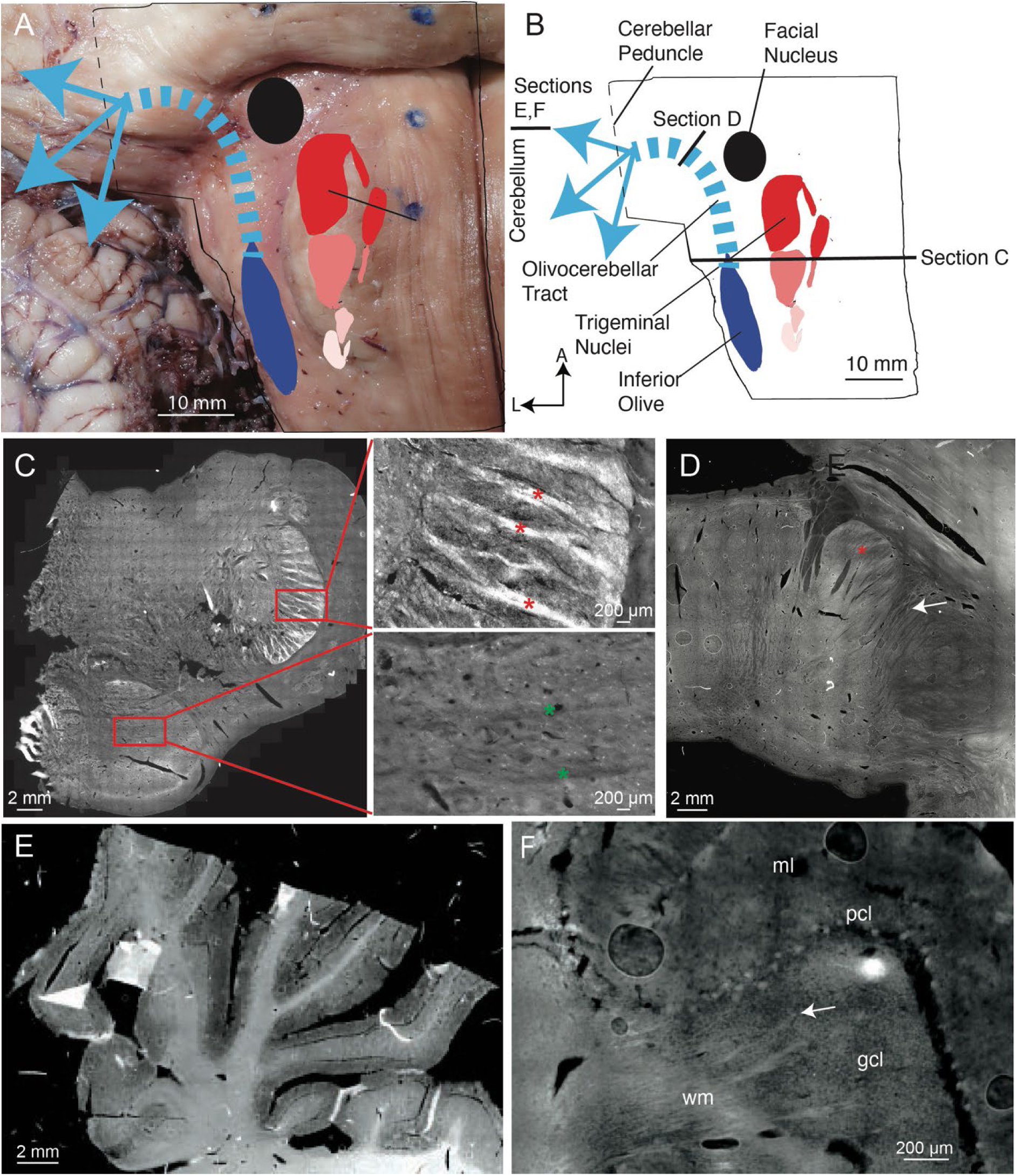
Peripherin-antibodies reveal climbing fibers and the olivo-cerebellar tract connecting the cerebellum to the inferior olive but not to the trigeminal nucleus. **A,** ventral view of the elephant brainstem with key structures schematically superimposed. **B,** schematic of the elephant brainstem. We indicate the positions of the peripherin-stained sections shown in C-F and our delineation of the olivo-cerebellar tract based on peripherin-reactive putative climbing fibers. **C,** left, overview picture of a brainstem section stained with anti-peripherin-antibodies (white color). Anti-peripherin-antibodies stain climbing fibers in a wide variety of mammals. The section comes from the posterior brainstem of African elephant cow Bibi; in this posterior region, both putative inferior olive and trigeminal nucleus are visible. Note the bright staining of the dorsolateral nucleus, the putative inferior olive according to Reveyaz et al. (the trigeminal nucleus according to Maseko et al., 2013). **C,** upper right, high magnification view of the dorsolateral nucleus (corresponding to the upper red rectangle in A). Anti-peripherin-positive axon bundles (red stars, putative climbing fibers) are seen in support of the inferior olive hypothesis of Reveyaz et al. **C,** lower right, high magnification view of the ventromedial nucleus (corresponding to the lower red rectangle in A). The ventromedial nucleus is weakly positive for peripherin but contains no anti-peripherin-positive axon bundles (i.e. no putative climbing fibers). Note that myelin stripes – green stars, weakly visible as dark omissions – are clearly anti-peripherin-negative. The region around the ventromedial nucleus is devoid of peripherin-reactivity and this is true throughout the brainstem. This observation suggests that the putative trigeminal nucleus does not receive climbing fiber input. **D,** anti-peripherin-antibody (bright color) stained section below the cerebellar peduncle. Putative climbing fibers (arrow) can be seen to be budding off the olivo-cerebellar (red star) tract into the cerebellar peduncle. **E,** anti-peripherin-antibody (bright color) stained section of the elephant cerebellum. The cerebellar white matter is bright, the cerebellar granule cell layer has a light grey appearance, and the cerebellar molecular layer has a dark grey appearance. Accordingly, peripherin-reactivity mirrors the distribution of cerebellar climbing fibers (white matter > granule cell layers > molecular layer). **F,** high magnification view of an anti-peripherin-antibody stained section of the elephant cerebellum. Climbing fibers are apparent by their elongated axonal pattern (arrow). Purkinje cells appear as bright dots. Even higher magnification revealed that the inside of Purkinje cells is dark (anti-peripherin-negative), i.e. the bright dot appearance of Purkinje cells reflects the ensheating of Purkinje cells by climbing fibers. wm = white matter; gcl = granule cell layer; pcl = Purkinje cell layer; ml = molecular layer.

### A delineation of the olivo-cerebellar tract supports our partitioning scheme

Previous work on the elephant brainstem (Shoshani et al., 2006; Maseko et al., 2013; Verhaart and Kramer, 1958; Verhaart 1962) suggested that the structure we assigned as trigeminal nucleus is in fact the inferior olive; this matter is also discussed in depth in our earlier discussions with the referees, which are published along with our article. We performed additional antibody staining to assess the possibility that the structure we assigned as trigeminal nucleus corresponds to the inferior olive. To differentiate between the trigeminal nuclei and the inferior olive, we used a climbing fiber antibody staining. Peripherin is a cytoskeletal protein that is found in peripheral nerves and climbing fibers. Specifically, climbing fibers of various species (mouse, rabbit, pig, cow, and human; Errante et al 1998) stain intensely with peripherin-antibodies. In Figure 2A we provide an overview of the elephant brainstem and key structures therein. Figure 2B shows schematically the key conclusions of our peripherin-antibody-staining, our delineation of the olivo-cerebellar tract, and indicates the position of peripherin-antibody-stained sections. We observed peripherin-reactivity in axonal bundles (i.e. in putative climbing fibers), in what we think is the inferior olive (Figure 2C, left and upper right, red stars). We also observe some peripherin-reactivity in what we think is the trigeminal nucleus, but not the distinct and strong labeling of axonal bundles (Figure 2C lower right, green stars). This lack of peripherin-reactive axon bundles suggests, that there are no climbing fibers, in what was previously thought of as the inferior olive of the elephants. We followed peripherin-reactive fibers through the brainstem and found they discharge into the cerebellar peduncle as expected for the olivo-cerebellar tract (Figure 2D, white arrow). Peripherin-reactivity was also observed in the cerebellum (Figure 2E), where putative climbing fibers ascend through the white matter and the granular cell layer and ensheath Purkinje cell somata (Figure 2F). These observations show that the strongly serrated dorsolateral nucleus connects to the cerebellum via climbing fibers. In contrast, the ventromedial nucleus, the putative trigeminal nucleus, does not receive climbing fibers. These data support our partitioning scheme over the assignments of Maseko et al 2013.

### Trigeminal nuclei in coronal and horizontal sections of African elephant brainstem

We found that putatively trunk-related trigeminal modules repeat at least two and probably four times in the anterior-posterior direction in the elephant brainstem. All these repeating modules had a higher cytochrome oxidase reactivity and a higher cell density than surrounding brainstem structures. Such repeats of trigeminal representations in the anterior-posterior direction are also seen in other mammals (Belford & Killackey, 1979; Ma, 1991). We refer to these modules with the same terminology as established in rodents. As in other mammals, we found the most anterior representation to be larger than the others, and we refer to this representation as nucleus principalis (Pr5, which stands for principal trigeminal nucleus). Figure 3A shows a Nissl-stained coronal section through the principalis trigeminal modules. In Figure 3B, C we provide a color-coded putative assignment of principalis modules. We assigned the large (grey) module to the trunk, because of its cytochrome oxidase reactivity, its elongation, and its extraordinary size. We assigned the elongated (red/pink) module to the nostril for the following reasons: 1. its unusually (among brainstem modules) thin tube-like appearance. 2. The widening towards the tip of the putative trunk module. 3. The cellular continuity with the mouth opening of the putative trunk module. 4. The topographic relationship with the lower jaw module, which matched the topography of the elephant head (Figure 3C). 5. The fact that this module had the same length as the trunk module. 6. The fact that there were no indications of a nostril module inside the putative trunk module, (where we initially expected a nostril representation). Our reasons for assigning the compact (blue) module to the lower jaw were its shape (Figure 3A-C) and topographic position. Next, we provide an overview of the arrangement of trigeminal modules in horizontal sections (Figure 3D-F, proceeding from dorsal to ventral). At the dorsal level (Figure 3D) only two trunk modules (TM), can be recognized. These are Pr5 and Sp5o, which stands for spinal trigeminal nucleus pars oralis TM, directly posterior to the Pr5. The cell density is low, the modules barely stand out from the surroundings and we think that at the dorsal level proximal trunk parts are represented. At the midlevel (Figure 3E) four repeating trunk modules (Pr5 TM, Sp5o TM, and Sp5i, which stands for spinal trigeminal nucleus pars interpolaris, and Sp5c, which stands for spinal trigeminal nucleus pars caudalis TM) can be recognized. We did not investigate facial representations other than the trunk module. The identification of the Sp5i and Sp5c trunk modules is only tentative at this point. The analysis of horizontal and parasagittal sections pointed to a mirror image-like arrangement of these modules. The cell density is higher at midlevel (Figure 3E). At the ventral level (Figure 3F) only two trunk modules (Pr5 and Sp5o TM, directly posterior to the principalis) can be recognized. In this section, the mirror image-like arrangement of the Pr5 TM and the Sp5o TM is evident. The cell density is very high and we think that the trunk tip is represented here. We conclude that repeating trigeminal trunk modules can be recognized in the elephant brainstem.

**Figure 3.**
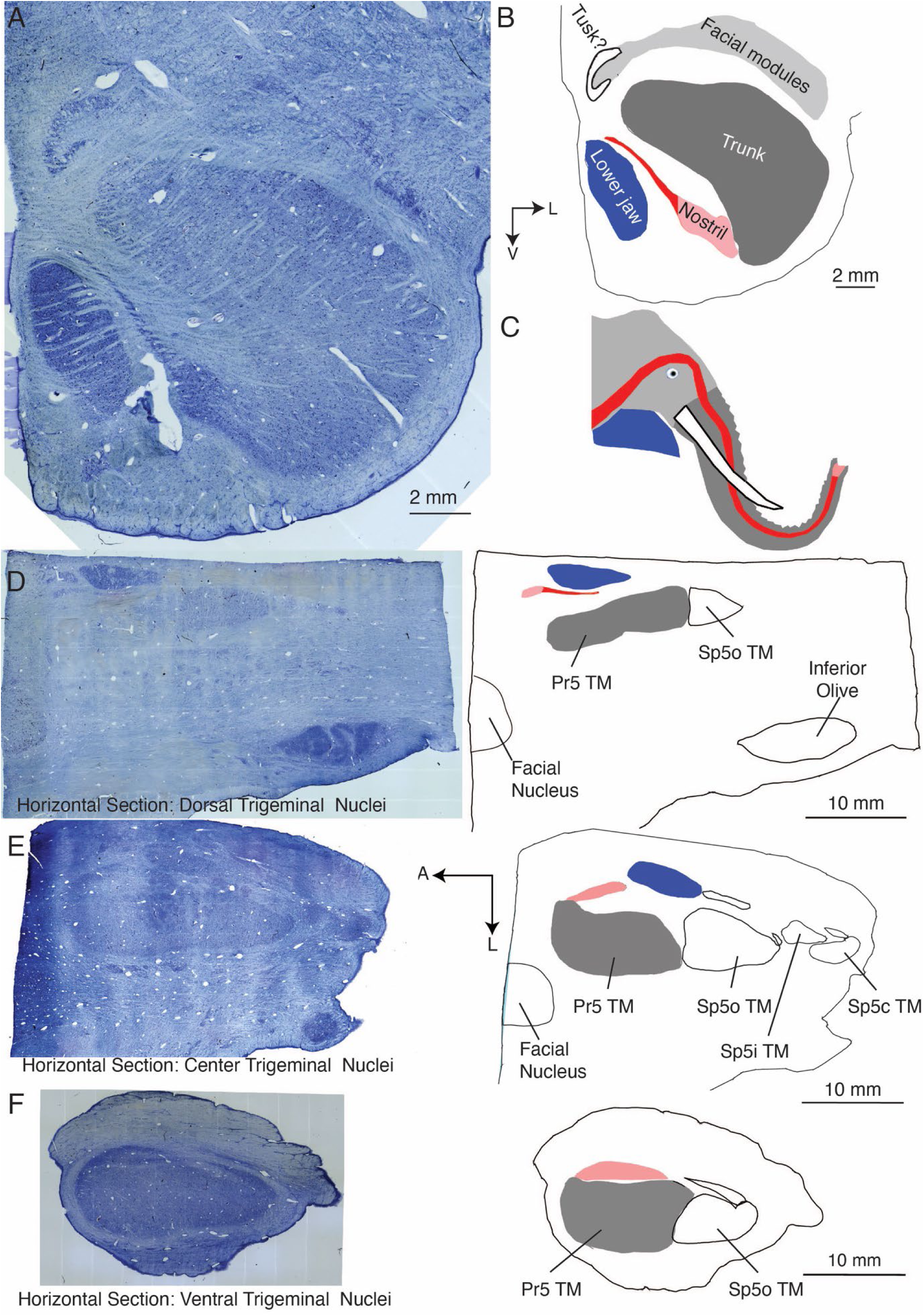
Overview of trigeminal nuclei in coronal and horizontal sections of African elephant cow Indra. **A**, micrograph of Nissl-stained coronal 60-µm-section through the right-hemispheric principalis trunk module of elephant cow Indra. The principalis is the most anterior and by far the largest trigeminal representation in elephants. **B**, drawing of color-coded trigeminal modules belonging to the principalis nucleus. **C**, drawing of an African elephant head with different facial structures color-coded according to the trigeminal modules they putatively correspond to in **A**, **B**. **D**, left micrograph of Nissl-stained horizontal section through the left-hemispheric trigeminal nuclei of elephant cow Indra. The section is positioned at a dorsal level of the trigeminal nuclei. Right, drawing of trigeminal modules shown in the left micrograph; for the principalis (Pr5) module the same color code as in **B**, and **C** has been used. Sp5o refers to the trunk module directly posterior to the principalis nucleus. The facial nucleus and the putative inferior olive were also identified. **E**, conventions as in **D**. Horizontal section through the mid-level of the trigeminal nuclei. Sp5i TM and Sp5c TM, refer to putative trunk modules posterior to the Sp5o TM. **F**, conventions as in **D**. Horizontal section through the ventral level of the trigeminal nuclei. Pr5, principal trigeminal nucleus; Sp5o, spinal trigeminal nucleus pars oralis; Sp5i, spinal trigeminal nucleus pars interpolaris; Sp5c, spinal trigeminal nucleus pars caudalis; TM = trunk module; A = anterior; L = lateral; V = ventral.

### The cellular architecture of the putative principalis trunk module

We studied the module’s cellular organization by Golgi stains of the putative principalis trunk module of the Asian elephant Raj (Figure 4A). Golgi stains identified two prominent neuronal elements. First, we observed bundles of large diameter (3-15 µm) axons, which run orthogonal to the module’s main axis in coronal sections (Figure 4B arrows). These axon bundles correspond to the myelin stripes described in Figure 5. Somato-dendritically stained neurons were the second neuronal element identified by Golgi stains (Figure 4C). The density of Golgi-stained neurons was very low. We reconstructed neurons using a Neurolucida system and superimposed 47 neurons (from three adjacent sections) in Figure 4D. The most abundant cells in the trigeminal nucleus were putative astrocytes (Figure 4E, green). We distinguished two types of neurons, putative principal neurons with large somata and branched dendrites (n = 41; black in Figure 4E) and other neurons, putative interneurons, with small somata and unbranched dendrites (n = 6; red in Figure 4E). Putative astrocytes, principal neurons, and interneurons differed markedly in their morphologies (Figure 4E, Table 1). The dendritic trees of neurons were elongated (Figure 4E), an observation confirmed when we prepared raw polar plots of dendritic orientation (Figure 4F, upper). We had the impression that dendritic elongation and axon bundles followed the same axis. We tested the idea that dendritic trees were aligned to myelin stripes by rotating dendritic trees and aligning all trees according to the local axon bundle orientations. When we aligned dendritic trees this way, we observed an even stronger population polarization of dendritic trees, i.e., dendritic trees were average twofold longer along the axon bundle axis (Figure 4F, lower).

**Figure 4.**
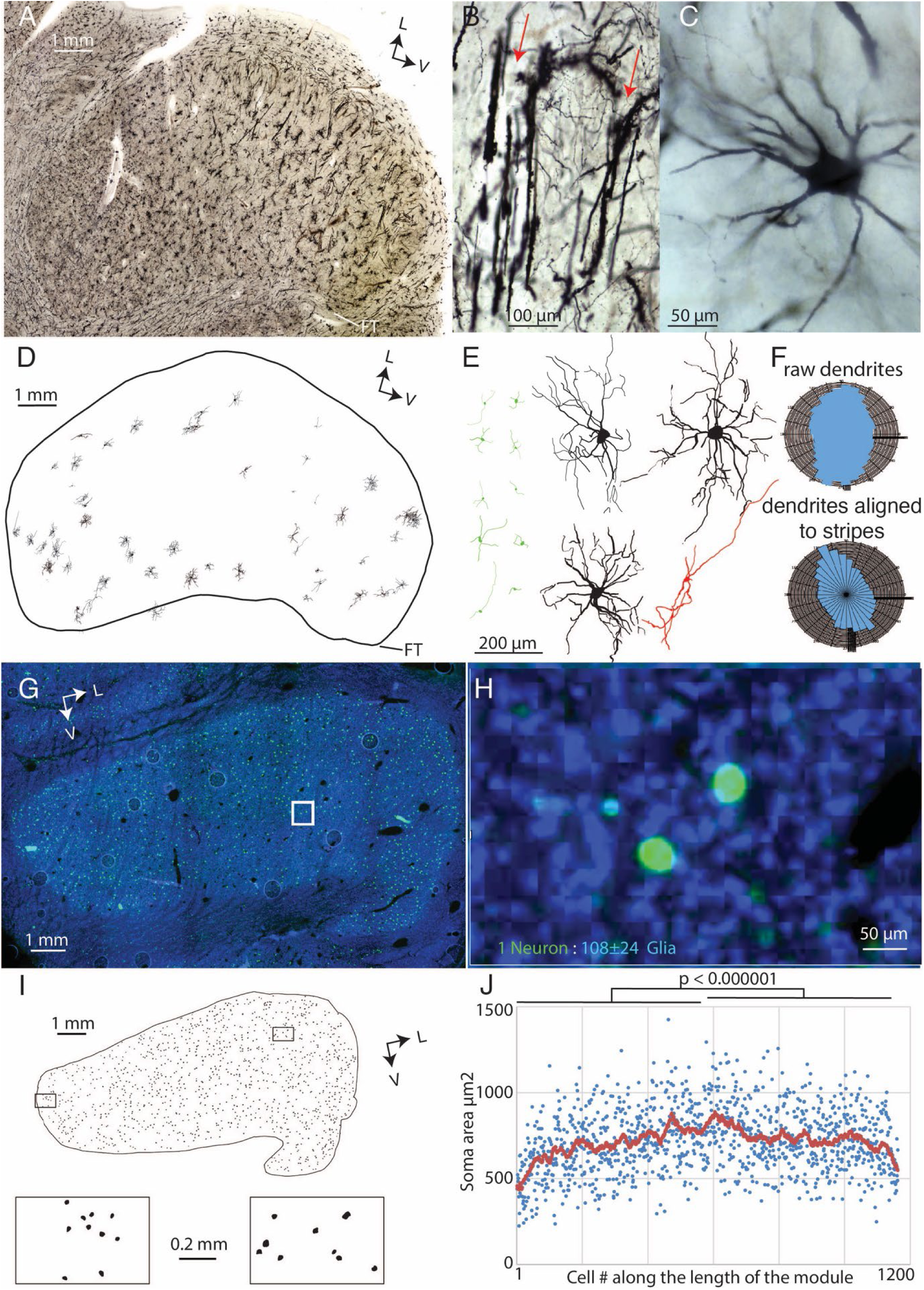
Cellular organization of the putative trunk module in the elephant trigeminal nuclei. **A**, Golgi stained 200 µm coronal section through the trunk module of Raj, a four-year-old Asian elephant bull. **B**, bundles of very thick axons are revealed by the Golgi staining (red arrows). Bundles were oriented orthogonal to the main axis of the module except for the putative finger representation, where they curved around. **C**, Golgi-stained neurons are also observed albeit at a low frequency. **D**, well-stained and well-preserved neurons reconstructed with a Neurolucida system. We show 47 neurons reconstructed from three adjacent coronal Golgi sections superimposed. Note the small size of the neurons relative to the module. **E**, left (green), ten putative astrocytes. These small cells are the most abundant cellular element in the trigeminal nucleus. Right, four neuronal reconstructions are shown at higher magnification. Putative principal cells (three shown in black, total n = 41) had large somata and branched dendrites. A few cells (one cell shown in red, total n = 6) had small somata and unbranched dendrites. Dendritic trees were weakly polarized. **F**, upper, raw polar plot of the orientation of neuronal dendritic segments (from all putative principal cells and putative interneurons, n = 47) relative to the soma confirms the common elongation of dendrites. Lower, when cells were aligned to the local axon bundle orientation an even stronger polarization of dendrites is evident. **G**, antibody staining of neurons (green fluorescence, NeuN-antibody) and nuclei of all cells (blue fluorescence, DAPI) of a coronal section through the putative trunk module of Indra, a 34-year-old female African elephant. Neuron density is low. **H**, high magnification view of the section shown in G, non-neural cells outnumber neurons by about a hundredfold (data refer to neuron and non-neural cell counts from three elephant trigeminal nuclei). **I**, upper, somata drawing from a Nissl stained 60 µm coronal section through the putative trunk module of Indra. Lower, cells from the medial, the putatively proximal trunk representation of the module, and the lateral, the putatively distal trunk representation. Note the soma size difference. **J**, plot of soma area along the length of the module. Neurons were sequentially measured along the axis of the module. Each dot refers to one of 1159 neurons in the section; red running average (across 40 neurons) of soma area. Cells are significantly larger in lateral (putatively distal trunk representation), unpaired T-test. FT, putative dorsal Finger Tip representation; V = ventral; L = lateral.

**Figure 5.**
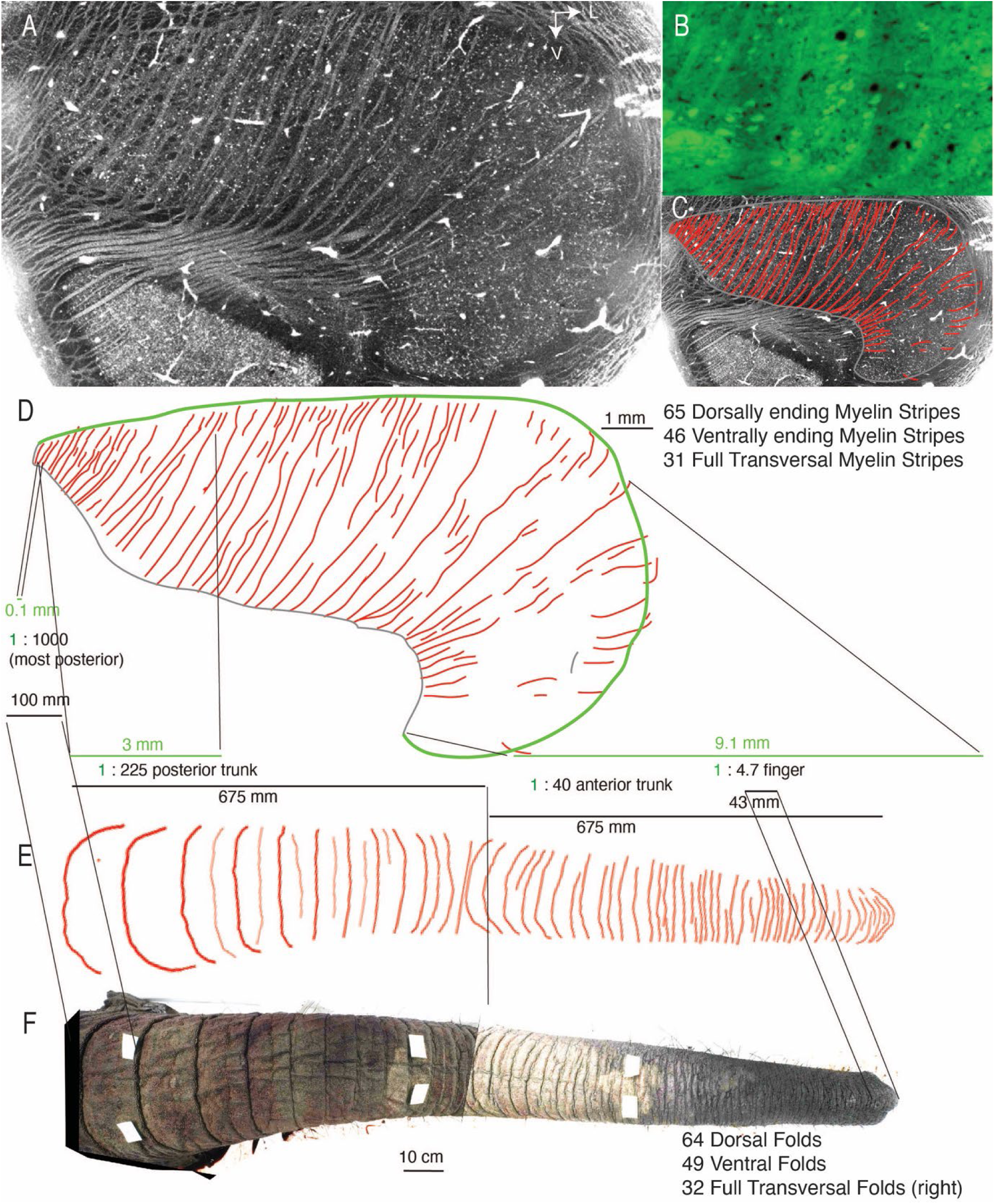
Trunk module myelin stripes form a precise map of trunk folds. **A**, a brightfield image of a freshly cut 60 µm coronal section through the center of the putative trunk module of adult female elephant Indra. Neurons are evident as small white dots and whitish myelin stripes are readily apparent even in this unstained tissue. **B**, a fluomyelin stain (green fluorescence) confirms that stripes contain myelin. High-resolution brightfield microscopy (not shown) and Golgi stains (Figure 2B) confirm that the stripes consist of axon bundles. **C**, a line drawing (red) of myelin stripes superimposed to the micrograph shown in **A**. **D**, upper, enlarged view of line drawing (red) of myelin stripes shown in **C**; we quantified dorsally ending, ventrally ending, and full transversal stripes. Such numbers match the number of trunk folds quantified in **E**. Lower, based on the idea of a match of trunk folds with myelin stripes one can compute magnification factors across the trunk module. Neural data (green) refer to distances between dorsally ending myelin stripes (i.e., neural distances were measured along the dorsal border (green line) of the module). Trunk distances were measured between dorsal trunk folds. **E**, drawing of the dorsal folds of the trunk of Indra. **F**, the composite photograph of the dorsal trunk of Indra. We counted dorsal folds, ventral folds (not visible here), and folds that fully transversed the right trunk side (not visible here). Trunk folds match in number, orientation (typically transversal), and patterning with myelin stripes seen on the trunk module. V = ventral; L = lateral.

**Table 1.** Morphological properties of elephant trigeminal cells in the Asian elephant Raj. Data (mean ± SD) comes from Golgi-stains soma diameter which was defined as the maximal Feret diameter. Data relate to n = 20 for putative astrocytes, n = 41 for putative principal neurons, and n = 6 for putative interneurons. In unpaired t-tests, all morphological parameters were significantly different between groups. See Figure 2.

| Cell type | Soma Diameter | Soma Area | # of Dendrites (Processes) | Dendritic (Process) Nodes | Total Dendritic (Process) Length |
| --- | --- | --- | --- | --- | --- |
| Putative Astrocytes | 8.5 ± 3.1 μm | 37 ± 25 μm<sup>2</sup> | 3.8 ± 1 | 1.1 ± 1 | 179 ± 99 μm |
| Putative Principal Neurons | 43 ± 9 μm | 1002 ± 420 μm<sup>2</sup> | 8.9 ± 2.9 | 22 ± 15 | 3344 ± 1783 μm |
| Putative Interneurons | 11.7 ± 4.9 μm | 85 ± 82 μm<sup>2</sup> | 5.8 ± 1 | 1.4 ± 1 | 551 ± 137 μm |

We stained coronal sections with NeuN-antibody to identify neurons and used the DNA-stain DAPI to identify all cell nuclei (Figure 4G). Neuronal density was fairly low, but the density of non-neuronal cells was substantial (Figure 4H). In counts of individual fluorescence sections, we observed a ratio of neurons to non-neural cells of 1 to 80 (in one elephant). We then made more systematic counts of neurons vs. visually identified non-neural cells across three entire trunk modules from two elephants. We then observed a ratio of 1 neuron to 108 ± 24 non-neural cells. We also observed that neuron size was not homogeneous across the putative trunk module. We observed a cell size increase from proximal to distal in all putative modules (n = 12), for which we had cellular stains. To quantify cell size differences, we drew somata from a Nissl-stained section through the center of the trunk module of the African elephant cow Indra (Figure 4I). We found cell size to increase significantly from the putative proximal to the putative distal finger representation (Figure 4J). We conclude that the putative trunk module contains transversally running axon bundles and neurons, which are vastly outnumbered by non-neural cells.

### Module myelin stripes match with number, orientation, and patterning of trunk folds

In Nissl or cytochrome-oxidase stains, we observed prominent myelin stripes apparent as white omissions. Remarkably, entirely unstained freshly-cut coronal brainstem sections showed the clearest stripe pattern in brightfield microscopy (Figure 5A). Fluorescent stains for myelin (fluomyelin) confirmed the presence of myelin (Figure 5B). As already suggested by Golgi staining, the myelin stripes appeared to consist of large-diameter axons. The visibility of myelin stripes varied with the sectioning plane and anterior-posterior position. Myelin stripes were most obvious at the anterior-posterior center of the putative trunk module, as seen for the coronal section in Figure 5A. We investigated the myelin-stripe trunk correspondence. To this end, we made drawings of myelin stripes (Figure 5C) and compared the pattern of myelin stripes (Figure 5D upper) to the pattern of trunk folds of Indra’s trunk (Figure 5E, F). Myelin stripe and trunk fold patterns were very similar. In all trunk modules sectioned, we observed an overall match of stripe and trunk fold orientation (to the module and trunk main axis, respectively). We also observed in all modules a lack of fully transversing stripes in the putative finger region of the module, which is consistent with the lack of folds across the trunk ‘mouth’. In favorable cases, where we had brightfield images of the trunk module and had access to the elephant’s trunk, the data hinted at a 1 to 1 matching of stripes and folds. Specifically, we observed 65 myelin stripes ending on the dorsal side of the module, 46 ventrally ending myelin stripes, and 31 full transversal myelin stripes. In terms of folds, we observed 64 dorsal trunk folds (Figure 5E), 49 folds on the ventral side of the trunk, and 32 folds that fully transversed the right side of Indra’s trunk. This numeric correspondence is very suggestive and inspired a detailed mapping of trunk sensory topography (Figure 5D lower). Based on stripe-wrinkle matching, we suggest that sensory magnification increases from 1000:1 (trunk: trigeminal nucleus) in the proximal representation of the trunk module to 5:1 in the trunk-finger representation. Sectioning angle was a major factor in determining the match between myelin stripes and folds, i.e. we observed myelin stripes in trunk fold-like patterns in all coronally sectioned specimens. In horizontally sectioned elephant brainstem, myelin stripes were seen but could not be related to folds. In parasagitally sectioned brainstems few myelin stripes were obvious. In coronal sections, myelin stripes were most obvious in the center of the module and matched best to trunk folds.

The staining method was another determinant of the match. Myelin stripes were best visible in unstained freshly cut sections with brightfield microscopy. As expected, myelin stripes were also stained positively for fluorescent myelin dyes, such as fluomyelin-green (Figure 5B) or fluomyelin-red (data not shown). Nissl or cytochrome oxidase stains were less sensitive than visualizing myelin stripes in brightfield images, i.e. not all stripes are visible in each section. While in some sections like the one shown in Figure 5D, pretty much all stripes could be mapped to trunk folds, most sections contained a few stripes that had deviating trajectories from the other stripes and these stripes could not be mapped to trunk folds. A good match of myelin stripes to folds depended also on the assessment of trunk folds. Specifically, a good match was only obtained, if we restricted fold counts to major trunk wrinkles/folds, minor trunk wrinkles appear not to be robustly represented by myelin stripes. We conclude myelin stripe patterns behave not unlike rodent cortical barrel patterns, the visibility of which also greatly depends on the staining method and sectioning angle.

### Myelin stripe architecture and the lack of a relation of stripes to trigeminal neurons

Their large size makes determining the architecture of myelin stripes challenging. To confront this challenge, we applied synchrotron-powered X-ray phase-contrast tomography of an 8 mm unstained and paraffin-embedded trigeminal nucleus tissue punch (Figure 6A, B); based on phase-contrast this methodology allows us to sample large image volumes (Figure 6C) with submicrometer (0.65 µm isotropic voxel size) resolution. Such imaging allowed us to identify myelin stripes in unstained trigeminal tissue (Figure 6D) and even enabled the reconstruction of individual large-diameter axons for several millimeters through the entire volume image (Figure 6E). As observed before with light microscopy, myelin stripes ran in the coronal plane and were about seven myelinated axons wide (maximum extent in the coronal plane; Figure 6F). Myelin stripes were circular axon bundles (Figure 6G) and were also about seven myelinated axons high (maximum extent in the anterior-posterior plane; Figure 6H). With that, we estimated that stripes are made up of 20-50 myelinated axons, unmyelinated axons could not be resolved in our analysis. The large-diameter axons, which could be followed through the X-ray tomography volume image followed an ‘all the way’ pattern (i.e. fully transversing the module). We also found that myelin stripes have a fairly consistent thickness from their dorsal to their ventral end (Figure 6I). This observation argues against the idea that myelin stripes are conventional axonal supply structures, from which axons divert off into the tissue. We also analyzed myelin stripes throughout the trunk module (Figure 6J) to understand how their thickness relates to trigeminal neuron numbers (i.e. the number of neurons between myelin stripes; Figure 6J). We observed little obvious relation between myelin stripe thickness and trigeminal neuron number. We conclude that myelin stripes have a stereotyped architecture, but show little relation to trigeminal neurons.

**Figure 6.**
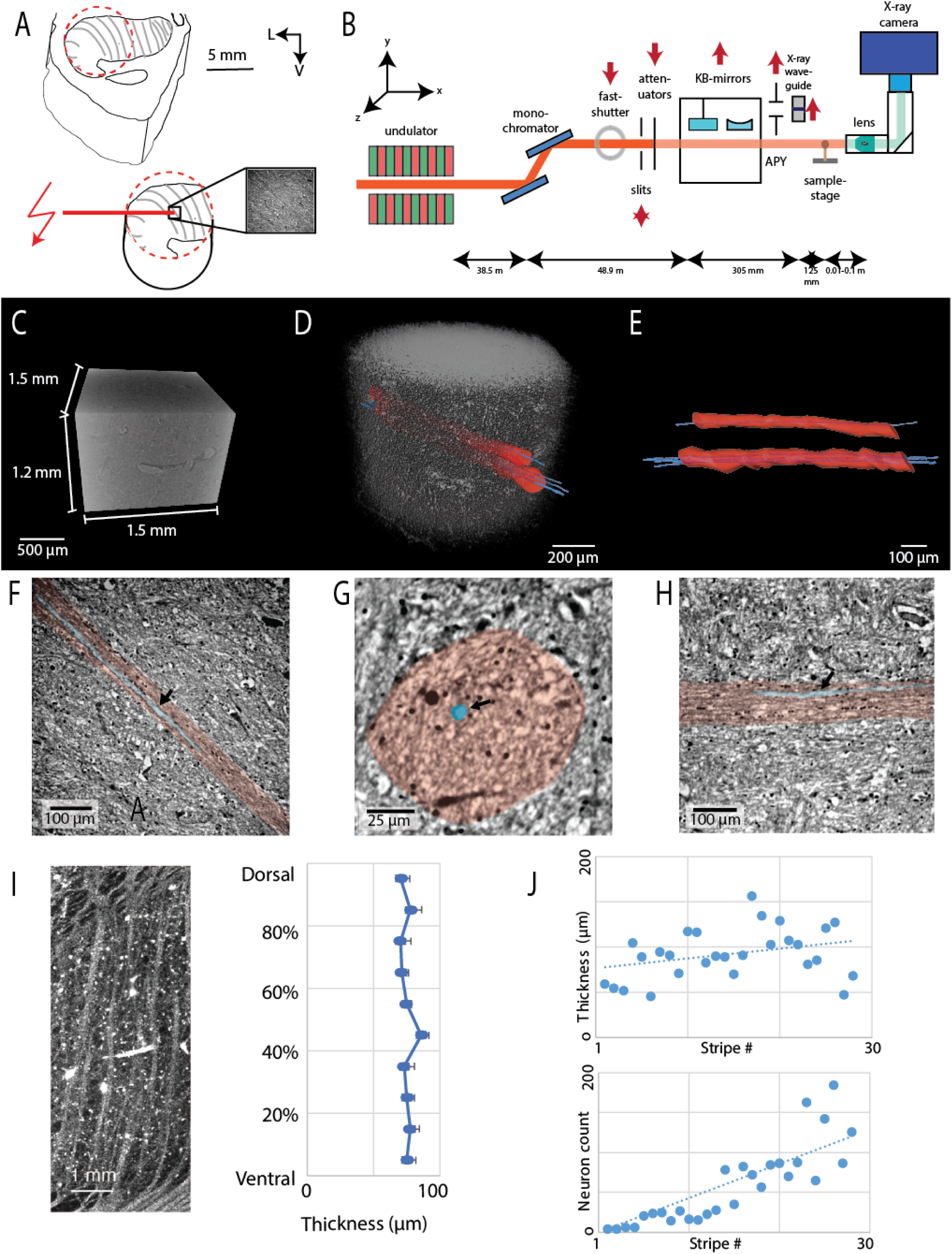
Microscopic organization of myelin stripes and absence of a strong relation of stripes to trigeminal neurons. **A,** upper, schematic of the trunk module with myelin stripes (grey) of African elephant Bambi and targeting of the 8 mm tissue punch (dashed red circle). Lower, synchrotron radiation (red flash, DESY, Hamburg) was directed to the area of the punch (black box) and imaged. V = ventral; L = lateral. **B**, sketch of the parallel beam setup of the GINIX endstation (P10 beamline, DESY, Hamburg). In this geometry, a dataset of the trunk module was acquired at an effective voxel size of 650nm3 **C**, dimensions of the imaged volume shown as a volume rendering (0.65 µm isotropic voxel size) **D**, transparent image volume. Two myelin stripes were followed through the volume image (highlighted red axon bundle). Four large diameter (∼ 15 µm) axons were also reconstructed and could be followed through the entire volume image (blue). **E**, axon bundles (red) and reconstructed axons (blue) in isolation. **F**, image section in the coronal plane at the center of the myelin stripe. The myelin stripe (pink overlay) is readily visible, the reconstructed axon is highlighted in blue, and the bundle has a width of about 7 myelinated axons. **G**, cross-section through the axon bundle (pink overlay). The virtual section is cut orthogonal to the coronal plane. The reconstructed axon is highlighted in blue. **H**, image section orthogonal to the coronal plane, the virtual section is cut in an anterior-posterior direction parallel to the axon bundle (pink overlay). The myelin stripe is readily visible, the reconstructed axon is highlighted in blue, and the bundle has a height of about 7 myelinated axons. **I**, left, myelin stripes. Right, measurement of the thickness (width orthogonal to the main stripes axis) along the dorsoventral axis of stripes; data are the average of ten measurements along ten myelin stripes that fully transversed the module. Stripe thickness is fairly constant, there is no evidence of stripe tapering as would be expected if axons bud off into the tissue. **J**, upper, measurement of the thickness (width orthogonal to the main stripes axis) of myelin stripes across the putative trunk module; only full transversal stripes were measured. Stripe thickness is fairly constant. Lower, neuron number between full transversal myelin stripes. Neuron number varies more than 100-fold and is very low (zero) between medial stripes (in the putatively proximal trunk representation). The fact that stripe thickness changes little across the module, while neuron number between stripes changes massively argues against a relationship between stripe thickness and trigeminal neuron number.

### The putative trunk module mirrors species differences in trunk folds and trunk use

We found that the trigeminal bumps on the ventral brainstem differ significantly between African and Asian elephants (Figure 7A). We counted neurons in the principalis trunk module and found that African elephants (740210 ± 51902, mean ± SD) had more neurons than Asian elephants (636447 ± 69729, mean ± SD) and also had a larger volume principalis trunk module (Figure 7B). Supplementary Tables 2 and 3 provide further information on our counts of the trigeminal nuclei. As noted there, the dorsal finger accounted for a large fraction (∼20%) of the trunk modules. We wondered, why brainstem bumps differed between African and Asian elephants, and therefore closely investigated the shape of trunk modules in these species. A cytochrome-oxidase stained coronal section through the trunk module of the African elephant Indra is shown in Figure 7C. Drawings of coronal sections from this trunk module and the trunk module of other African elephants are shown in Figure 7D; myelin stripes (violet) were visible as whitish omissions of the cytochrome oxidase or of Nissl stains. We also determined the length, the width, and the longitudinal position of the greatest width of the module (black line; Figure 7D). A cytochrome-oxidase-stained coronal section through the trunk module of Asian elephant Raj is shown in Figure 7E and drawings from this and other trunk modules of Asian elephants are shown in Figure 7F. Basic aspects of the module trunk were similar in African and Asian elephants (Figure 7C-F), but the details differed. First, the African elephant trunk modules had fewer and thicker myelin stripes (Figure 7D, F); this is a most interesting observation since African elephants have fewer trunk folds than Asian elephants. Asian elephants have more of these folds, but they are also more shallow and less pronounced (Schultz et al., 2023). Given our limited material and that myelin stripes are less conspicuous in Asian elephants than in African elephants, we were not able to ascertain the one-to-one match of stripes and folds that we could show for the African elephant Indra in Figure 5. Second, African elephant trunk modules were significantly longer but not wider (Figure 7G). Asian elephant trunk modules had a much more round-shaped appearance than African elephant trunk modules. The greatest width of the Asian elephant trunk module is at positions representing the trunk wrapping zone, which we determined from photographs. African elephant trunk modules were also significantly more ‘top-heavy’, i.e. they had their widest point much closer to the putative trunk tip (Figure 7H). We suggest that the shape differences between African and Asian elephant trunk modules might be related to the different grasping strategies of these two elephant species (Racine, 1980). African elephants have two fingers and tend to pinch objects (Figure 7I; upper), a grasping strategy that emphasizes the trunk tip in line with their ‘top-heavy’ trunk module. Asian elephants in contrast have only one finger and tend to wrap objects with their trunk (Racine, 1980; Figure 5I; lower). This grasping strategy engages more of the trunk and, in line with this behavior, the width of the Asian elephant trigeminal nucleus is maximal in the trunk wrapping area.

**Figure 7.**
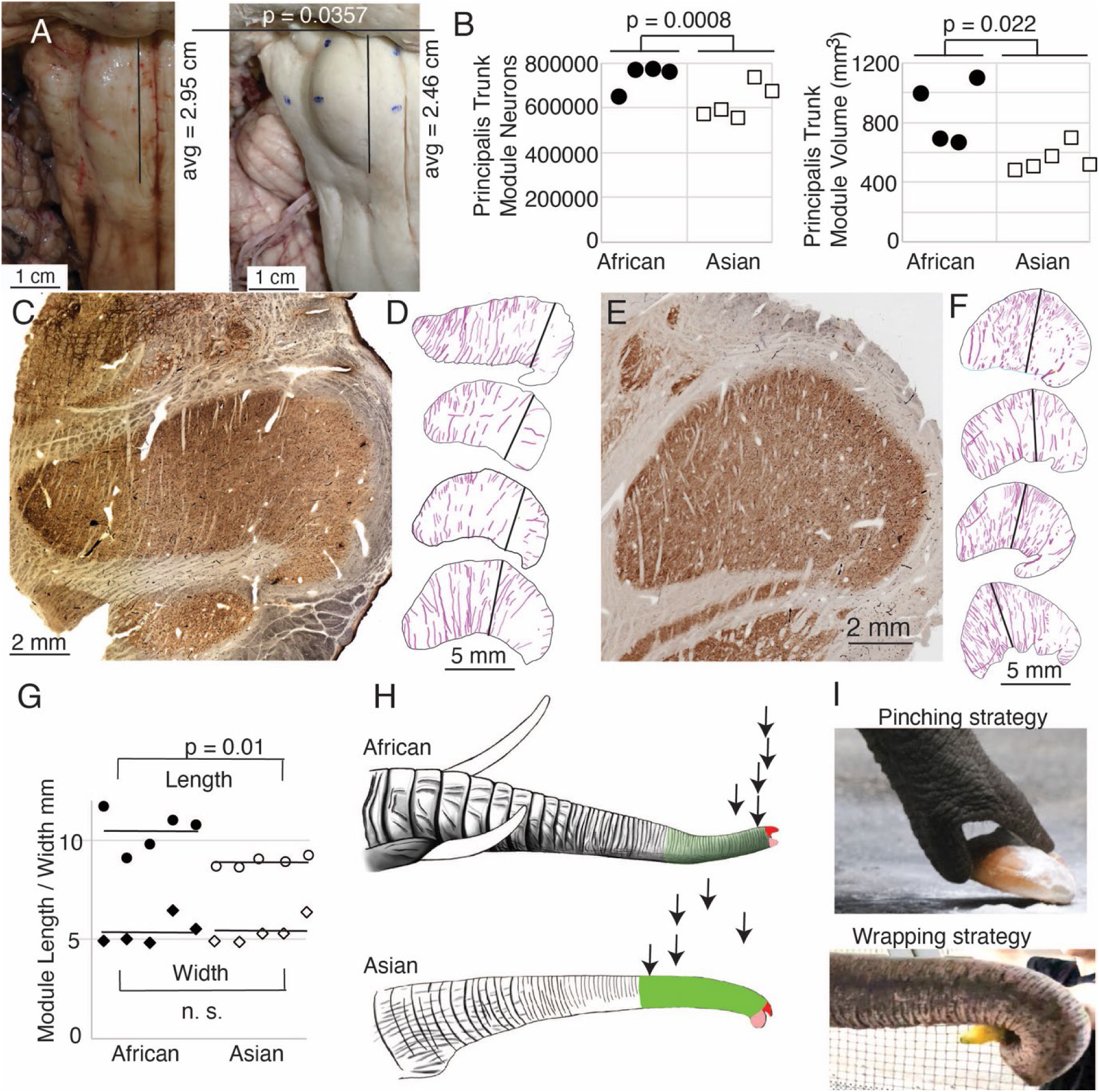
Differences between the putative trigeminal trunk modules of Asian (*Elephas maximus*) and African (*Loxodonta africana*) elephants. **A**, left, ventral view of the brainstem of African elephant Indra. Right, ventral view of the brainstem of Asian elephant Dumba. The anterior-posterior length of the trigeminal nuclei from the pons is indicated as a black line. The average trigeminal nuclei length refers to 10 African and 10 Asian trigeminal nuclei, the p-value refers to a Mann-Whitney test. The trigeminal nuclei bump is more elongated in African than in Asian elephants. **B**, neuron number (left) and volume (right) of the putative principalis trunk module in Asian and African elephants. Trigeminal nuclei come from three African and three Asian elephants; p-values refer to unpaired t-tests. **C**, micrograph of a cytochrome-oxidase-stained section through the putative trunk module of African elephant Indra. **D**, drawings of the outlines and myelin stripes from cytochrome oxidase or Nissl stained sections through the putative trunk module of African elephants; the top drawing was made from the micrograph shown in **C**. The black line refers to the point of greatest width along the direction of myelin stripes on the putative trunk shaft (the putative finger was not considered in the width analysis). **E**, micrograph of a cytochrome-oxidase-stained section through the putative trunk module of Asian elephant Raj. **F**, drawings, of putative trunk modules from Asian elephants; conventions as in **D**. **G**, length, and width of putative trunk module in African and Asian elephants. p-values refer to t-tests. **H**, upper, drawing of the trunk of an African elephant. Lower, drawing of the trunk of an Asian elephant; note that Asian elephants have more folds. Arrows mark the point of greatest width of the putative trunk module (black lines in **D, F**) projected back on trunk positions in African (upper) and Asian (lower) elephant trunks. We also highlighted in color the dorsal (red) and ventral (pink) trunk tip and wrapping zone of Asian elephants in green and the analogous trunk part of African elephants in light green. The extent of the trunk wrapping zone was determined from photographs of Asian elephants wrapping objects. Specifically, we defined the wrapping zone as the trunk parts in contact with large objects (mangos, melons, fodder beets) being wrapped. **I**, object grasping/pinching behavior in African (upper) and object wrapping strategy in Asian (lower) elephants (adapted from Kaufmann et al., 2022).

## Discussion

### Summary

We describe a pair of large bumps on the ventral surface of the elephant medulla that contain metabolically highly active, densely vascularized repeating modules. The trunk module contains an accurate myelin map of trunk folds. Mapping myelin stripes to the trunk folds indicated an increase in sensory magnification from the proximal to the distal trunk. Magnification analysis also identified an enlarged trunk wrapping zone in Asian elephants, who wrap objects with their trunk.

### The ventral brainstem bumps likely correspond to elephant trigeminal nuclei

Establishing elephant brainstem organization is challenging because both tracing methods and *in vivo* electrophysiology cannot be applied to elephants. Our assignments of trigeminal nuclei deviate from earlier suggestions (Shoshani et al., 2006; Maseko et al., 2013; Verhaart and Kramer, 1958; Verhaart 1962), which assigned the putative trigeminal nuclei as inferior olive, and the structure identified as inferior olive, as trigeminal nuclei. For several reasons, we think that our partitioning scheme with ventromedial trigeminal nuclei and a dorsolateral inferior olive is superior to the scheme of Maseko et al 2013 with a ventromedial inferior olive and dorsolateral trigeminal nuclei. Our synopsis of the evidence is the following. First of all, we agree with that concerning brainstem position our scheme of a ventromedial trigeminal nucleus and a dorsolateral inferior olive deviates from the usual mammalian position of these nuclei (i.e. a dorsolateral trigeminal nucleus and a ventromedial inferior olive). However, cytoarchitectonics support our partitioning scheme. The compact cellular appearance of our ventromedial trigeminal nucleus is characteristic of trigeminal nuclei. The serrated appearance of our dorsolateral inferior olive is characteristic of the mammalian inferior olive. To our knowledge, nobody has described a mammalian trigeminal nucleus with a serrated appearance, that would apply to the elephant according to Maseko et al. 2013. Furthermore, metabolic staining (cyto-chrome-oxidase reactivity) supports our partitioning scheme. Specifically, our ventromedial trigeminal nucleus shows intense cyto-chrome-oxidase reactivity as it is seen in the trigeminal nuclei of trigeminal tactile experts. Additionally, the myelin stripes on the ventromedial trigeminal nucleus are isomorphic to trunk wrinkles. Isomorphism is a characteristic of somatosensory brain structures (barrel, barrelettes, nose-stripes, etc) and we know of no case, where such isomorphism was misleading. The large-scale organization of the ventromedial trigeminal nuclei in anterior-posterior repeats is characteristic of the mammalian trigeminal nuclei. To our knowledge, no such organization has ever been reported for the inferior olive. Finally, the connectivity analysis supports our partitioning scheme. According to our delineation of the elephant olivo-cerebellar tract, our dorsolateral inferior olive is connected via peripherin-positive climbing fibers to the cerebellum. In contrast, our ventromedial trigeminal nucleus (the referee’s inferior olive) is not connected via climbing fibers to the cerebellum. The above arguments suggest that the elephant trigeminal nuclei very likely correspond to the ventral brainstem bumps (please also see our commentary to referees published along with the article). As clearly expressed by the comments of Referee 2 published along with our article, the strength of other partitioning schemes, which assign the ventral brainstem bump as inferior olive is the better positional match of the assignment with the inferior olive of other mammals. We think this is a valid criticism. Our model offers coherent anatomical entities with trigeminal nuclei that look like trigeminal nuclei of other mammals, a trunk module with a striking resemblance to the trunk, and an inferior olive that looks like the inferior olive of other mammals.

### Detailed neuroanatomic mapping of elephant trigeminal nuclei

The protruding of the ventral brainstem bumps in the elephant brain reminds us of the protruding of layer 2 cell clusters in the human entorhinal cortex (Solodkin and Van Hoesen 1996). More generally speaking myelin stripes of the putative elephant trigeminal nuclei, are another example of isomorphic representation in the somatosensory system. Other classic examples of such isomorphic representation are cortical barrels (Woolsey and Van der Loos), brainstem barrelettes (Belford & Killackey, 1979) in the whisker system. Also nose-related isomorphisms have been described before, i.e. the pig cortical rostrum gyrus (Ritter al. 2021), or the stripe-like representation of nose appendages in the brainstem of the star-nosed mole (Catania, Leitch and Gauthier 2011). Our work provided detailed mapping of the elephant trigeminal brainstem into four repeating nuclei, consisting of several facial modules (most prominently the trunk module, the nostril module, and the lower jaw module). Because of the level of detail of our topography suggestions, however, we think it will be relatively straightforward to test the validity of our topography suggestions. Specifically, we predict that the dorsal trunk finger representation of the principalis trunk module will be connected with the distal part of the dorsal subnucleus of the facial nucleus (which contains the putative motor representation of the dorsal trunk finger (Kaufmann et al., 2022)). We would also predict that the dorsal trunk finger representation of the principalis trunk module will be connected with the dorsal trunk finger representation of the oralis nucleus. These connectivity suggestions are in the 1-2 cm range and can be tested with postmortem tracers like DiI. We also predict fewer myelin stripes in trunk modules of elephants with particularly few trunk folds (newborn or fetuses of African elephants) compared to adult African elephants.

### A myelin map of trunk folds, white matter function and myeloarchitecture

According to conventional wisdom, neurons (gray matter) are the site of processing, and myelinated axons (white matter) are a subjugated supply system, whose sole function is to bring the correct axonal input to neurons. Our observations on trunk module myelin stripes are at odds with this view of myelin. Specifically, myelin stripes show no tapering (which we would expect if axons divert off into the tissue). More than that, there is no correlation between myelin stripe thickness (which presumably correlates with axon numbers) and trigeminal module neuron numbers. Thus, there are numerous myelinated axons, where we observe few or no trigeminal neurons. These observations are incompatible with the idea that myelin stripes form an axonal ‘supply’ system or that their prime function is to connect neurons. What do myelin stripe axons do, if they do not connect neurons? We suggest that myelin stripes serve to separate rather than connect neurons. Specifically, trunk module myelin stripes look like a map of trunk folds. Myelin stripes match with the number, orientation, and species-specific patterning of trunk folds. We note that if myelin stripes would behave as an axonal ‘supply’ system, they would be very thin/invisible in the proximal trunk, proximal trunk folds would not be visible, and distal stripes should be very thick. If myelinated axons have a life of their own and do not simply go where they find target neurons, we need to analyze them in novel ways. In particular, it seems to be a good idea to ‘look’ at patterns of myelination, rather than to immediately assume that this is a supply/connectivity system. We note early neuroanatomists like Oskar Vogt (Vogt, 1911; Niewenhuys, Broere & Cerliani, 2015) described incredibly intricate patterns of intracortical myeloarchitecture, patterns that are not easily explained in terms of a connectivity system to this day. We reckon that the exciting novel methodologies for determining myelo-architecture (Haenelt et al. 2023) will bring the issues and unanswered questions raised here to the foreground of neuroscientific inquiry. In conclusion, we propose a novel white-matter function, which is to separate and functionally demarcate neurons as opposed to the conventionally assumed white-matter function of connecting neurons.

### Trigeminal organization in Asian and African elephants

At first sight, Asian and African elephant trigeminal nuclei are very similar. Both elephants have big ventral brainstem bumps, which contain the same modules (a putative trunk, nostril, and lower jaw module) and the nuclei stain intensely for cytochrome oxidase reactivity. A closer look reveals species differences, however, which may relate to the different trunk grasping strategies of Asian and African elephants. The first difference refers to the shape of the ventral brainstem bump, which is more roundish and shorter in Asian elephants and more elongated in African elephants. To our knowledge, this ventral brainstem bump is the only hitherto described difference, which allows us to differentiate Asian and African elephant brains from the outside. The length difference between Asian and African elephant ventral bumps reflects the different shapes of Asian and African elephant trunk modules. The African elephant trunk module is notably longer, more slender, and more top-heavy than the swaged Asian elephant trunk module. As we pointed out in Figure 7, such differences imply an enlargement of the trunk wrapping zone in the Asian elephant trunk module, in line with the object-wrapping behavior of Asian elephants (Racine 1980). Similarly, the top-heavy shape of the African elephant trunk module could be instrumental in the object-pinching of African elephants (Racine, 1980). These trigeminal differences are reminiscent of similar Asian-African species differences in the elephant facial nucleus (Kaufmann et al., 2022). We conclude that grasping behavior shapes the species-specific architecture of the trigeminal nuclei.

### Conclusion

The elephant brainstem is exquisitely well-ordered and contains very large and detailed trigeminal representations. Trunk module myelin stripes form a map of trunk folds and accordingly serve to functionally separate neurons rather than to connect them. Further work should test the predictions of sensory topographies outlined here and ask, what further insights the elephant brain provides about the organization of gray and white matter.

## Acknowledgments

Supported by BCCN Berlin, Humboldt-Universität zu Berlin and the Deutsche Forschungsgemeinschaft (DFG, German Research Foundation) under Germanýs Excellence Strategy – EXC-2049 – 390688087. We thank Andreea Neukirchner, Susanne Holtze, Guido Fritsch, Francisca Egelhofer, Aniston Sebastiampillai, Uta Westerhüs, Anne Nesseler, Karin Risse and Jana Petzold. Dr. Claudia Szentiks, Zoltan Mezö, Marc Gölkel and Katharina Brehm helped with necropsy. Several zoological institutions contributed, in particular the Berlin Zoo (Germany) and the Zoo Schönbrunn Vienna (Austria), as well as Zoo Augsburg (Germany), Opel-Zoo Kronberg (Germany), Zoo Poznan (Poland), Tierpark Hagenbeck (Germany), the Elefantenhof Platschow (Germany).

We acknowledge DESY (Hamburg, Germany), a member of the Helmholtz Association HGF, for the provision of experimental facilities. Parts of this research were carried out at PETRA III and we would like to thank Fabian Westermeyer for assistance in using the P10 GINIX endstation. Beamtime was allocated for proposal I-20220980.

## Materials and Methods

Our methods were described in detail in our recent publications (Kaufmann et al., 2022; Purkart et al., 2022) and we only repeat key aspects here.

### Elephant specimens

All specimens came from zoo elephants and were collected by the Leibniz-IZW (Leibniz Institute for Zoo and Wildlife Research, Berlin) over the last three decades in agreement with CITES regulations. All animals included in the study died of natural causes or were euthanized by experienced zoo veterinarians for humanitarian reasons, because of insurmountable health complications. An overview of the elephant specimen used in this study is provided in Supplementary Table 1.

*Asian elephants, Elephas maximus.* Data from four-year-old elephant bull Raj (Tierpark Hagenbeck, Germany), from the adult Asian elephant cow Burma (52 years old, Zoo Augsburg, Germany), and from the Asian elephant cow Dumba (44 years old, elephant farm Platschow, Germany). Different data were derived from the various Asian elephant specimens.

*African savanna elephants, Loxodonta africana.* Data from four adult African elephant cows: Zimba (39 years old, Opel-Zoo Kronberg, Germany), the 34-year-old elephant cow Indra (Platschow), and Bambi (38 years old, Hungary). Different data were derived from the various African elephant specimens.

*Specimen condition.* Specimen conditions varied widely in our study (for details see Kaufmann et al., 2022). Some heads or other material reached us frozen and none of the elephant heads/brains were perfused. Even though many of the animals included were dissected by professional veterinarians, the preservation of material varied across specimens. A variety of reasons contribute to the suboptimal preservation of elephant material. Specifically, it often takes days to dissect elephants and the animals’ carcasses cool down only very slowly. Furthermore, the freezing leads to freezing artifacts, and even in extracted brains fixative action is slow, because of elephant brain size. Some of these problems are discussed and have been solved (Shoshani, 1982; Manger et al., 2009).

### Elephant preparation and trigeminal nucleus collection

*Elephant preparation.* In adult elephants, heads and trunks were removed at the respective zoos and the remaining skull was trimmed with motorized saws and axes at the Leibniz-IZW Berlin. Some of the brains from trimmed skulls of adult elephants were extracted by Francisca Egelhofer and Aniston Sebastiampillai at the Neuropathology of the Charité, Berlin.

*Trigeminal nucleus extraction.* We proceeded with trigeminal nucleus collection after extraction of the brain and dura removal followed by several weeks of fixation in 4% paraformaldehyde solution. To remove trigeminal nuclei, we positioned entire elephant brains with their ventral side up in a dissection tray. We then dissected away blood vessels and the pia arachnoidea from the elephant brain stem. To dissect out trigeminal nuclei we oriented ourselves at the trigeminal nuclei bump shown in Figure 1B.

### Trigeminal nucleus sectioning, preparation, and staining

Trigeminal nuclei were stained for Nissl-substance. Most trigeminal nuclei were sectioned in 60 µm thickness with our cryotome. A series of sections were processed, alternating with Nissl and antibody staining (NeuN antibodies). We also performed Golgi and cytochrome oxidase reactivity (Wong & Kaas, 2008; Wong-Riley, 1979). The antibody staining procedure followed the protocols described by Purkart et al., 2022 and Kaufmann et al., 2022. For Golgi staining, brains were only minimally fixated (1 day 1% paraformaldehyde in 0.1 M phosphate buffer). Staining was performed with a commercial kit (Rapid Golgi Kit, Gentaur, Aachen Germany). Sections for Golgi staining were cut at a thickness of 200 µm. We additionally performed an antibody staining with anti-Peripherin antibodies (anti-Peripherin Antibody; AB 1530; Sigma – Aldrich) on sections with the putative trigeminal nucleus and inferior olive, following Purkart et al., 2022 and Kaufmann et al., 2022 protocol.

### Cellular measurements, somata drawings, and neuronal reconstructions

Thin Nissl-stained sections were viewed with Stereo Investigator software (MBF Bioscience, Williston, USA) employing an Olympus BX51 microscope (Olympus, Japan) with an MBFCX9000 camera (MBF Bioscience, Williston, USA) mounted on the microscope. The microscope was equipped with a motorized stage (LUDL Electronics, Hawthorne, USA) and a z-encoder (Heidenhain, Schaumburg, USA). Stereo Investigator software was used for stereological procedures, cell size, and axon diameter measurement and for acquiring images. Drawings of neural somata were also generated from Nissl-stained sections on this system. Digitized images were adjusted for brightness and contrast using Adobe Photoshop (Adobe Systems Inc., San Jose, Calif., USA), but they were not otherwise altered.

Neuronal reconstructions were prepared from Golgi stains on a Neurolucida system (Microbrightfield, USA).

### Stereology based on the optical fractionator

We used an optical-fractionator approach to quantify cell numbers in the trigeminal nuclei. An overview of the results and counting parameters used in our study is provided in Supplementary Table 2. Here estimated the total number with Stereo Investigator software (MBF Bioscience, Williston, USA) using a sampling scheme called the optical-fractionator method. Our region of interest was identified and outlined at low (2x objective) magnifications. The neurons were identified by their shape staining intensity and large size at high magnification (20x) and counted individually. Without exception, the trigeminal trunk module was well-defined by a higher neuron density than the surrounding brain structures. The standard stereological sampling scheme is independent of volume, measurements, and shrinkage because the number of neurons is estimated directly without referring to neuron densities. Using the optical-fractionator technique, we counted the nucleoli that came into focus and fell within the acceptance lines of the dissector, which were randomly placed on the series of sections (Kaufmann et al., 2022).

We counted neurons in the Nissl stains of 9 trigeminal nuclei of 6 elephants. We used the following parameters. The dissector laid a grid of squares over our region of interest with a size of 2000 x 1000 μm², where we counted the neurons at each dissector in the counting frame area of 350 x 350 μm^2.^ At each counting frame, we counted between 0 and 15 neurons. Around 1000 neurons were counted in each trigeminal nucleus to assess the total number of neurons (see Supplementary Table 2). The entire elephant trigeminal nucleus spanned ∼400 60-µm-sections in adult animals, every 20^th^ section was counted. The guard zone was set to zero. The mean thickness measured at every counting site was measured to be around 18 µm and used to estimate the total number of neurons.

### Paraffin embedding for X-ray phase-contrast tomography

A 2 x 2 x 2 cm³ sized trigeminal brainstem piece of an African elephant Bambi was immersed in an ascending ethanol series of 20/50/70% (1 d each) at 4 °C one week before paraffin embedding. Subsequently, the sample was infiltrated by first acetone, then xylol, and finally paraffin in an automatized vacuum paraffin infiltration processor. After cooling and hardening of the paraffin embedded sample overnight we obtained an 8 mm biopsy punch from the putative finger region of the trigeminal brainstem region.

### Synchrotron X-ray tomography

X-ray phase contrast volumes of the unstained and paraffin-embedded trigeminal nucleus were scanned with an unfocused, quasi-parallel synchrotron beam (PB) at the GINIX endstation, at a photon energy E_ph_ of 13.8 keV, selected by a Si(111) monochromator. Projections were recorded by a microscope detection system (Optique Peter, France) with a 50-m-thick LuAG: Ce scintillator and a 10× magnifying microscope objective onto a sCMOS sensor (pco. edge 5.5, PCO, Germany) (Frohn et al., 2020). This configuration enables a field-of-view (FOV) of 1.6 mm × 1.4 mm, sampled at a pixel size of 650 nm. The continuous scan mode of the setup allows the acquisition of a tomographic recording with 3000 projections over 360° in less than 2 min. Afterward, dark field and flat field images were acquired.

### Phase retrieval and tomographic reconstruction

First, the raw detector images were corrected by dark subtraction and empty beam division. In addition, hot pixel and detector sensitivity variations were removed by local median filtering. A local ring removal was applied around areas where wavefront distortions from upstream window materials did not perfectly cancel out after empty beam division. Phase retrieval was performed for each projection, using the linear CTF approach (Cloetens et al., 1999; Turner et al., 2004), implemented in the HoloTomoToolbox (Lohse et al., 2020). This implementation allows both for formulation of additional constraints as well as a nonlinear with iterative minimization of a Tikhonov-functional starting from the CTF result as an initial guess. However, for the unstained samples shown here, this was not found to be necessary. Apart from phase retrieval, the HoloTomoToolbox provides auxiliary functions, which help to refine the Fresnel number or to identify the tilt and shift of the axis of rotation (Lohse et al., 2020). Tomographic reconstruction of the datasets was performed by the ASTRA toolbox (van Aarle et al., 2016; van Aarle et al., 2015, using the iradon-function and a Ram-Lak filter.

### Volume image segmentation

Tomographic images were segmented in an extended version of the Amira software (AmiraZIBEdition 2022.17, Zuse Institute Berlin, Germany). A combination of the ‘lasso’ and ‘brush’ tools was used to manually label the axons and myelin stripes within the volume image. Labels were placed every 5 – 50 images and interpolated in between.

### Statistical analysis

All statistical tests are specified in the respective Figures, legends, or in the text. All tests were two-tailed.

## Supplementary Material

**Supplementary Table 1.** Overview of elephants and treatment of the corresponding specimen.

| **Name**<br>**(species)** | **Sex** | **Location at**<br>**death** | **Age (y),**<br>**died on**<br>**(dd/mm/yyyy)** | **Specimen**<br>**treatment** |
| --- | --- | --- | --- | --- |
| Zimba<br>(*Loxodonta*<br>*africana*) | F | Opel-Zoo<br>Kronberg,<br>Germany | 39, 10/04/2021 | Brain removed,<br>briefly frozen,<br>put in fixative |
| Indra<br>(*Loxodonta*<br>*africana*) | F | Elefantenhof<br>Platschow,<br>Germany | 34, 2022 | Brain removed,<br>put in fixative |
| Bambi<br>(*Loxodonta*<br>*africana*) | F | Circus, Hungary | 38, 2023 | Brain removed,<br>put in fixative |
| Dumba<br>(*Elephas*<br>*maximus*) | F | Elefantenhof<br>Platschow,<br>Germany | 44, 18/02/2022 | Brain removed,<br>put in fixative |
| Burma<br>(*Elephas*<br>*maximus*) | F | Zoo Augsburg,<br>Germany | 52, 16/06/2021 | Brain removed,<br>put in fixative |
| Raj (*Elephas*<br>*maximus*) | M | Tierpark<br>Hagenbeck,<br>Germany | 4, 2022 | Brain removed,<br>put in fixative |
| Bibi (*Elephas*<br>*maximus*) | F | Zoo Aalborg,<br>Denmark | 41, 2023 | Brain removed,<br>put in fixative |

**Supplementary Table 2.** Optical fractionator counts of the principalis trunk module. Cell count estimates were derived using the optical fractionator; every twentieth section was sampled. See text for details.

| Specimen | Cells sampled | Gundersen Coefficient of error | Principalis Trunk Module Neuron Estimate |
| --- | --- | --- | --- |
| Indra left Principalis (African) | 965 | 0.03 | 650690 |
| Bambi right Principalis (African) | 986 | 0.03 | 762349 |
| Zimba left Principalis (African) | 995 | 0.05 | 773112 |
| Zimba right Principalis (African) | 993 | 0.04 | 774690 |
| **Average African Elephants (± SD)** | **985** | **0.0375** | **740210 ± 51902** |
| Burma left Principalis (Asian) | 857 | 0.04 | 738813 |
| Burma right Principalis (Asian) | 792 | 0.04 | 677816 |
| Dumba left Principalis (Asian) | 1291 | 0.03 | 571419 |
| Dumba right Principalis (Asian) | 1347 | 0.03 | 592418 |
| Raj left Principalis (Asian) | 853 | 0.04 | 556728 |
| **Average Asian Elephants (± SD)** | **1028** | **0.036** | **636448 ± 69730** |

**Supplementary Table 3.** Optical fractionator counts for other trigeminal modules. Cell-count estimates were derived using the optical fractionator; every twentieth section was sampled. See text for details.

| Structure | n | Cells sampled | Gundersen Coefficient of error | Neuron Estimate |
| --- | --- | --- | --- | --- |
| Nostril Module<br>Average African Elephants (± SD) | 4 | 90±12 | 0.11 ± 0.01 | 70247±12888 |
| Nostril Module<br>Average Asian Elephants (± SD) | 4 | 101±33 | 0.11 ± 0.02 | 61046±13748 |
| Dorsal Finger<br>(Part of Principalis Module)<br>Average African Elephants (± SD) | 4 | 234±34 | 0.0875±0.025 | 163799±29470 |
| Dorsal Finger<br>(Part of Principalis Module)<br>Average Asian Elephants (± SD) | 4 | 196±74 | 0.1±0.03 | 119919±23722 |
| Sp5o, spinal trigeminal nucleus pars oralis Trunk Module<br>Average African Elephants (± SD) | 3 | 342±45 | 0.07±0.01 | 296464±55463 |
| Sp5o, spinal trigeminal nucleus pars oralis Trunk Module<br>Average African Elephants (± SD) | 5 | 401±34 | 0.05±0.005 | 273910±29700 |

